# Single-cell transcriptomics uncovers that cell-to-cell communication controls p53 activation

**DOI:** 10.1101/2025.05.30.657016

**Authors:** Leandro Castellano, Mark E Robinson, Aleksandra Dabrowska, Niklas Feldhahn, Silvia Ottaviani, Justin Stebbing

## Abstract

Conventional chemotherapy remains a cornerstone of cancer treatment and is highly effective in the adjuvant and neoadjuvant settings. However, therapeutic resistance, especially in the metastatic context, limits its long-term efficacy. While significant progress has been made in elucidating mechanisms of chemotherapy resistance, the role of cell-to-cell communication in modulating treatment response remains largely unexplored. Here, we combined microfluidic technology with single-cell RNA sequencing (scRNA-seq) to directly compare the transcriptional responses of cells in communication versus those in isolation. Upon dox treatment, we observed robust activation of the p53 pathway in communicating cells, whereas this response was markedly impaired in isolated cells. Moreover, pharmacological inhibition of caveolar-mediated endocytosis mimics the loss of p53 activation, further implicating this pathway in the communication-dependent regulation of chemotherapy response. This finding uncovers a previously unrecognized and essential role for paracrine signalling in mediating p53-dependent responses to chemotherapy. Our study highlights the critical importance of cell-to-cell communication in shaping therapeutic outcomes and opens new avenues for understanding and potentially overcoming chemotherapy resistance.

## Introduction

Chemotherapy is a key therapeutic approach for the treatment of many types of cancer^1^. It exploits the vulnerabilities of rapidly dividing cancer cells by damaging their DNA, interfering with DNA replication, and ultimately leading to cell death^2^. However, the DNA damage response (DDR) either activates repair mechanisms or leads to cell death when the damage is irreparable, which is a frequent outcome in cancer cells with defective DDR pathways^2^.

A critical player in this process is the tumour suppressor protein p53, a transcription factor that orchestrates the DDR^3^. Upon activation, p53 can induce cell cycle arrest, providing time for DNA repair. If the damage is too extensive, p53 can trigger apoptosis, promoting cell death or senescence, where damaged cells stop dividing but remain metabolically active^3^. P53 selectively transcribes genes belonging to different pathways, determining cells’ fate in response to this stress^3^.

In the absence of DNA damage p53 is continuously degraded by the E3 ubiquitin ligase MDM2^4^. When stress occurs, p53 is phosphorylated rendering it resistant to MDM2^5^. This mechanism allows a rapid increase of p53 when stress ensues^3,6^. The DDR has recently been characterised at the single-cell level using live-cell reporters^7,8^ and single-cell RNA sequencing (scRNA-seq)^9^. These studies have provided valuable insights into how p53 and its targets, including CDKN1A, regulate the choice of pathways that determine cell fate.

Chemotherapy resistance presents a significant challenge to effective cancer treatment, highlighting the importance of understanding the mechanisms underlying this process—an area of intense ongoing research^10^. Under chemotherapy, a subset of tumour cells can survive the treatment^11^. Intriguingly, single-cell live imaging studies investigating p53 dynamics have shown that, following genotoxic insults in HCT116 colon cancer cell lines, the rate of p53 accumulation determines whether the cells survive the treatment^7^. Differential p53 activation was also found to drive varying levels of expression of genes in the inhibitor of apoptosis (IAP) family, thereby promoting resistance to chemotherapy^7^.

However, to the best of our knowledge, the importance of cell-to-cell communication through paracrine signalling for cell fate and p53 activity in response to genotoxic stress has never been investigated. This is crucial, as cancer cells may detach from the primary tumour, invade and form single circulating tumour cells no longer regulated by paracrine signalling from their original microenvironment. These cells are transported to other organs via the bloodstream, leading to metastasis^12^. It is essential to determine whether single tumour cells released from the tumour bulk lacking paracrine communication remain sensitive to chemotherapy or develop resistance. Understanding this aspect can help create new therapeutic strategies to prevent metastasis.

To investigate this aspect, we assessed cell fate following treatment with the genotoxic chemotherapy drug doxorubicin (dox) in individual cells isolated within sealed microfluidic chambers (on-chip, OCS), combined with single-cell transcriptomics. By preventing cell-to-cell communication, dox treatment failed to activate the p53 pathways, highlighting an indispensable yet previously unrecognised role of paracrine signalling in the DDR response.

## Results

### scRNA-seq reveals distinct temporal signatures of dox treatment

To dissect the DDR at a single-cell time series during a timecourse, we stimulated HCT116 cells with the DNA damaging agent dox for 0, 4, 8 and 24 hours (h) and performed scRNA-seq using the Fluidigm C_1_ microfluidics platform. We used one C_1_ chip (96 capture sites) per time point. We monitored capture efficiency by microscope imaging and excluded doublets and debris from the analysis. Cells were filtered for minimum feature coverage and fraction of reads mapping to mitochondrial or ERCC spike-in control features. After stringent quality criteria, we successfully profiled 157 cells (44 cells vehicle, 33 cells dox 4h, 40 cells dox 8h and 40 cells dox 24h) for this experiment. We sequenced filtered cells to an average depth of 3.3(±1.2) million reads per cell, covering 7.5(±1.2) thousand features. Next, we applied Remove Unwanted Variation (RUV) normalization to remove batch effects by covariate detection between control samples. To define a reference timeseries of response to dox, we performed pseudotemporal trajectory inference on bulk-treated cells using SCORPIUS. Unbiased clustering analysis revealed distinct signatures for each timepoint. Importantly, these clusters were not associated with a specific cell cycle phase as assigned by a pre-trained marker pairs classifier (**Fig. 1A**). Key DNA-damage response genes such as CDKN1A and MDM2 exhibited high expression variability in unstimulated cells but were coherently up-regulated following dox treatment, especially at late timepoints (**Fig. S1**). This demonstrates that the DNA damage response occurs uniformly across stimulated cells. Pseudotemporal analysis of the single cells identified a primary trajectory of response (**Fig. 1B**). Interestingly, by analysing single-cell gene expression profiles we were able to identify 4 main gene expression modules (M1-4), that were contributing to the pseudotemporal trajectory (**Fig. 1C**). M1 (n=170) was significantly associated with early repressed genes, M2 (n=145) with late repressed genes, M3 (n=101) with early activated genes and M4 (n=80) with late activated gene sets (**Fig. 1C**). Further we revealed the top enriched pathways in each module (**Fig. 1D**) together with the top 10 genes contributing to the trajectory in each module (**Fig. 1E**). While repressed genes (M1 and M2) were particularly enriched for transcripts coding for factors that modulate mitochondrial function and core transcription and translation machinery, activated genes (M3 and M4) were enriched for genes involved in the DNA damage response, cell cycle, and p53 activity, as expected.

**Figure 1.**
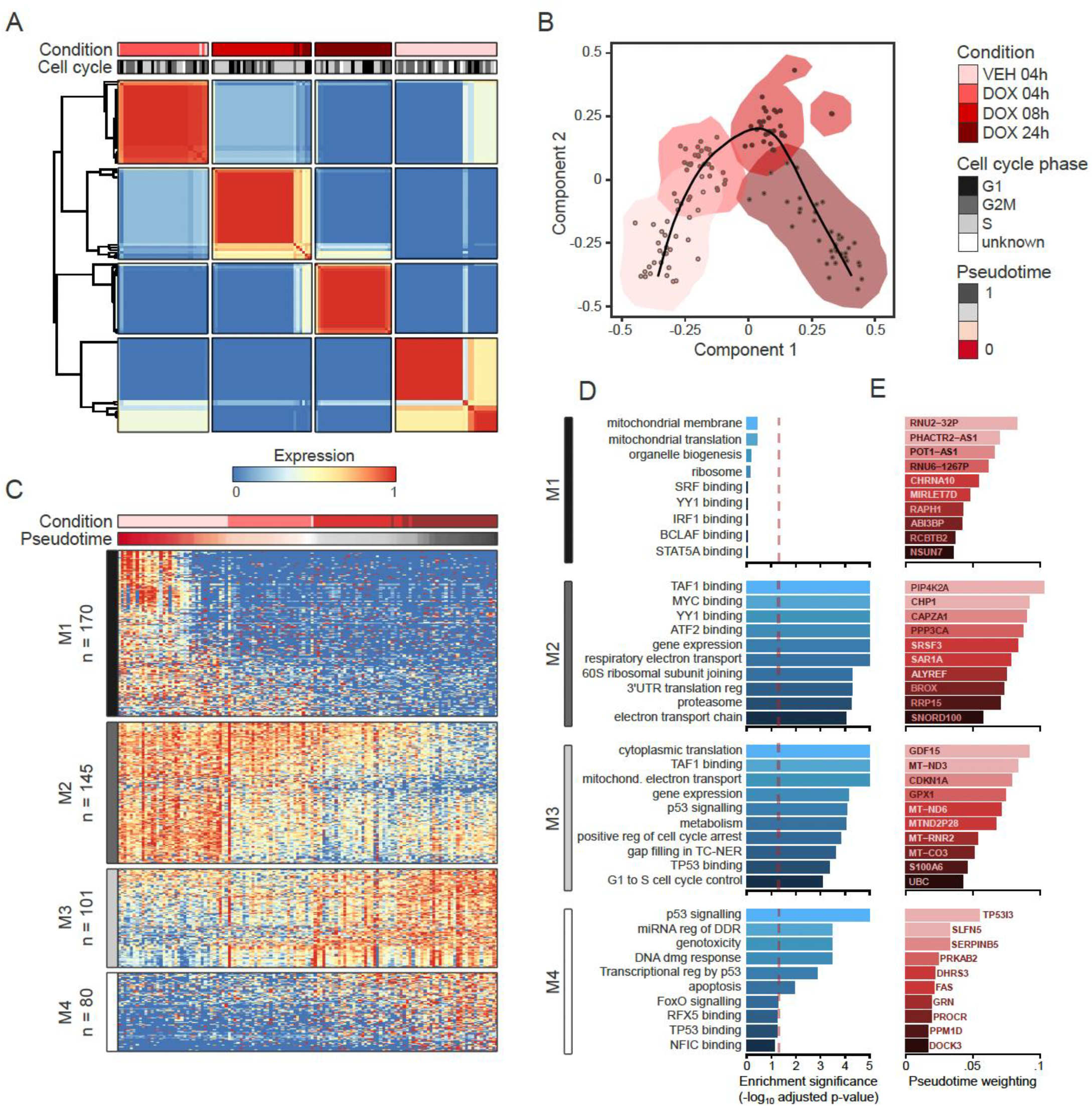
Response to doxorubicin treatment in single cell timeseries. **(A)** Unsupervised clustering of cells reveals distinct signatures of doxorubicin treatment (DOX) at each timepoint compared to vehicle control (VEH) and indicates no strong bias in cell cycle stage. **(B)** Pseudotemporal ordering of single cells identifies the primary trajectory of response. **(C)** Four main modules of genes contributing to the pseudotemproal trajectory were identified (M1-4) corresponding to early repressed, late repressed, early activated and late activated gene sets. **(D)** Top 10 enriched pathways in each module. **(E)** Top 10 genes contributing to the trajectory in each module.

### The DDR is impaired by lack of cell-to-cell communication

To evaluate whether the DDR induced by dox was influenced by cell-to-cell communication, we applied a modified C_1_ workflow developed by Shalek and colleagues^13^. According to this protocol, cells were captured in the C_1_ chip, treated with media containing dox or vehicle, and then immediately sealed in individual chambers to prevent intercellular communication for 4 hours (on-chip stimulation, OCS). Following incubation, samples were processed for scRNA-seq as for the previous timecourse experiment. Before we could compare the differential gene expression analysis of dox between the OCS and the 4h treatment from the time course experiment [bulk (BLK) method], two key confounding factors needed to be addressed. When cells were treated with dox or vehicle in the timecourse experiment, cells grew in adherent conditions and in the presence of serum. However, once the cells are individually captured on the C_1_ chip they are in a low adherent surface and in the absence of serum. To address these confounding factors and ensure that only the lack of paracrine factors was evaluated, we performed C_1_ single-cell RNA-seq on four additional conditions. We grew HCT116 cells in low adherent conditions with or without serum and with dox or vehicle for 4 hours. After stringent quality criteria, we successfully profiled 80 cells in the OCS (40 cells vehicle, 40 cells dox 4h), and 165 cells in the low adherent conditions (40 cells vehicle with serum, 42 cells dox with serum, 49 cells vehicle without serum and 34 cells dox without serum). Filtered cells were sequenced to an average depth of 3.8(±1.9) million reads per cell, covering 6.8(±1.3)k features, average mitochondrial fraction 15.1(±9.6)%; adherence, serum, or treatment method had no consistent effect on cell quality (**Fig. S1A**). All filtered cells were integrated and RUV-normalized to remove unwanted batch variance (**Fig. S1B**) without removing the effects of treatment and on-chip stimulation (**Fig. 2A**). Normalized cells cluster along 2 primary axes corresponding to treatment timepoint and methodology (on-chip vs bulk). Difference in response to dox (treatment) in OCS versus BLK (method) was assessed by modelling interaction effect (treatment:method), while controlling for confounding effects (serum and adherence). Differential gene expression analysis was performed between all conditions showing the effect of confounding factors (**Fig. 2B-C)**. Ultimately the interaction effect condition (OCS_DOC_int) shows the effect that lack of paracrine signalling has on gene expression in response to dox after eliminating all confounding variables. Surprisingly, blocking cell-to-cell communication showed impairment of typical p53 regulated processes such as apoptosis pathway, repression of MYC targets and loss of P53 pathway response in OCS response compared to BLK by GSEA analysis (**Fig. 2C**).

**Figure 2.**
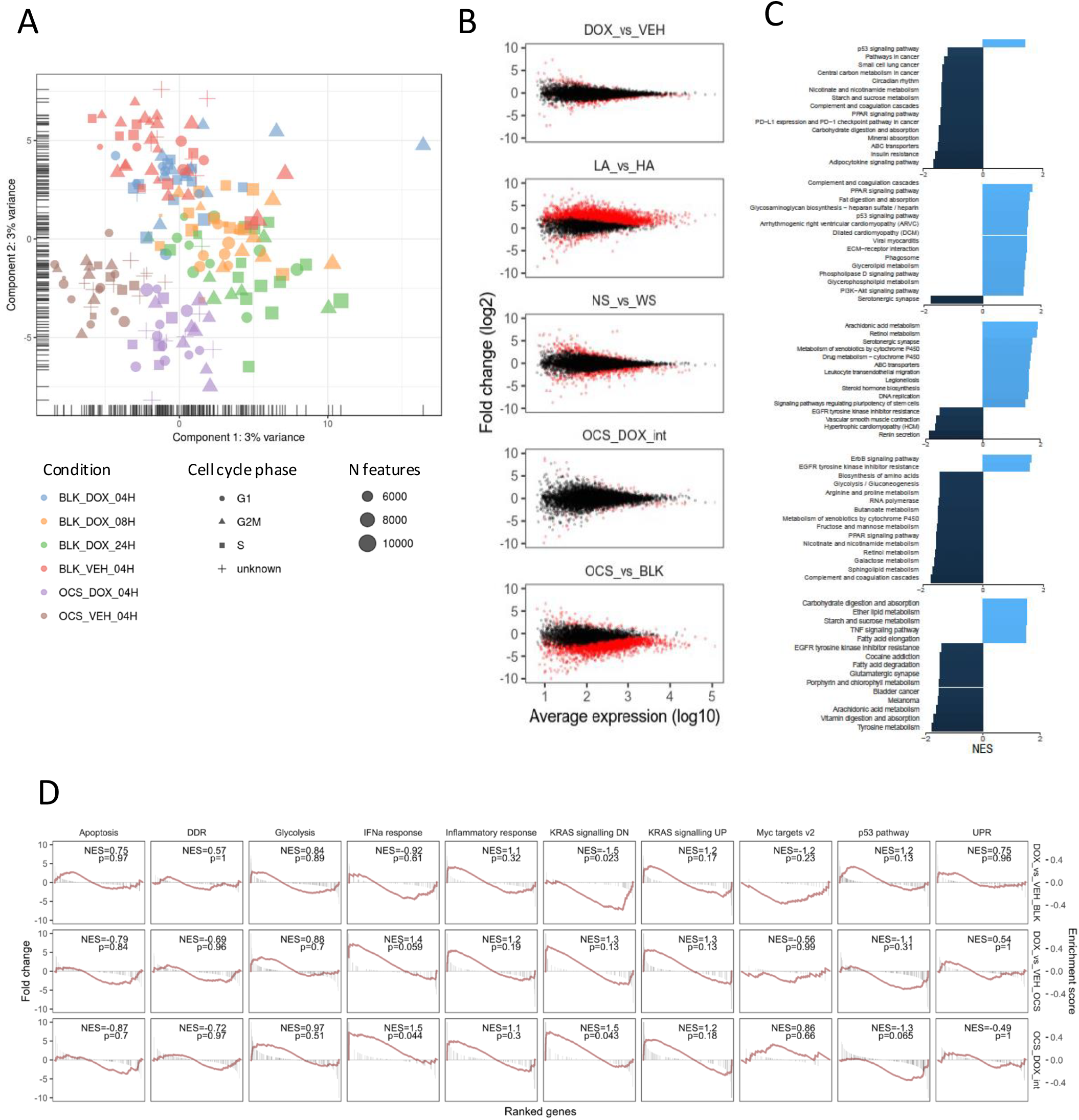
Effect of doxorubicin stimulation after experimentally blocking cell-to-cell communication. **(A)** PCA of integrated dataset - normalization removes major batch effects without eliminating effects of treatment or on-chip stimulation. **(B)** MA plots for overall differentially expressed genes in each comparison. **(C)** Gene set enrichment for each comparison. **(D)** GSEA shows loss of apoptosis pathway, loss of repression of MYC targets and loss of P53 pathway response in on chip response compared to bulk. *Abbreviations:* DOX=doxorubicin, VEH=vehicle, LA=low adherent condition, HA= adherent condition, NS=no serum, WS=whole serum, OCS= on-chip stimulation, int= interaction effect, BLK=bulk method.

### Absence of cell-to-cell communication impairs p53 signalling

The unexpected loss of p53-regulated signalling in cells stimulated with dox but lacking cell-to-cell communication prompted us to investigate this further. We built an expression map combined with p53 pathway analysis of p53-related genes by directly comparing the expression fold change of each gene in BLK (left panel) and OCS (right panel) (**Fig. 3**). Several p53-regulated genes were strongly downregulated in OCS conditions including p14^ARF^/CDKN2A, MDM2, p21/CDKN1A, GADD45 suggesting an overall inhibition of p53 response in non-communicating cells. This was further confirmed by analysing the expression of key p53 response genes in BLK compared to OCS conditions upon dox treatment (**Fig. S2**). As expected, dox induced upregulation of p53 response genes compared to vehicle in both adherent and serum and low adherent and no serum conditions, however, no upregulation was seen in OCS (**Fig. S2**). To further investigate this, we performed Ingenuity Pathway Analysis (IPA) comparison analysis (**Fig. S3**). This analysis validated our previous finding showing that p53 signalling together with DNA damage, and mitochondrial dysfunction were no longer statistically enriched in OCS condition. Interestingly, endocytosis related signalling were less enriched in OCS condition compared to BLK. These were the caveolar-mediated endocytosis signalling and clathrin-mediated endocytosis signalling, suggesting that these pathways could play a role in impairing p53 signalling in OCS condition. To test this hypothesis further, we treated cells with Genistein (an inhibitor of the caveolar-mediated endocytosis signalling) for 10 minutes followed by treatment with vehicle or dox (4h) and measured p53 signalling activation via gene expression of p53-responsive genes (**Fig. 4A**). Inhibition of caveolar-mediated endocytosis signalling significantly impaired the upregulation of p53-responsive genes induced by dox, suggesting that this pathway is crucial for the activation of p53 response. However, when we inhibited the clathrin-mediated endocytosis using PitStop2, no effect was seen, indicating that the clathrin-mediated endocytosis was not involved (**Fig. 4B)**.

**Figure 3.**
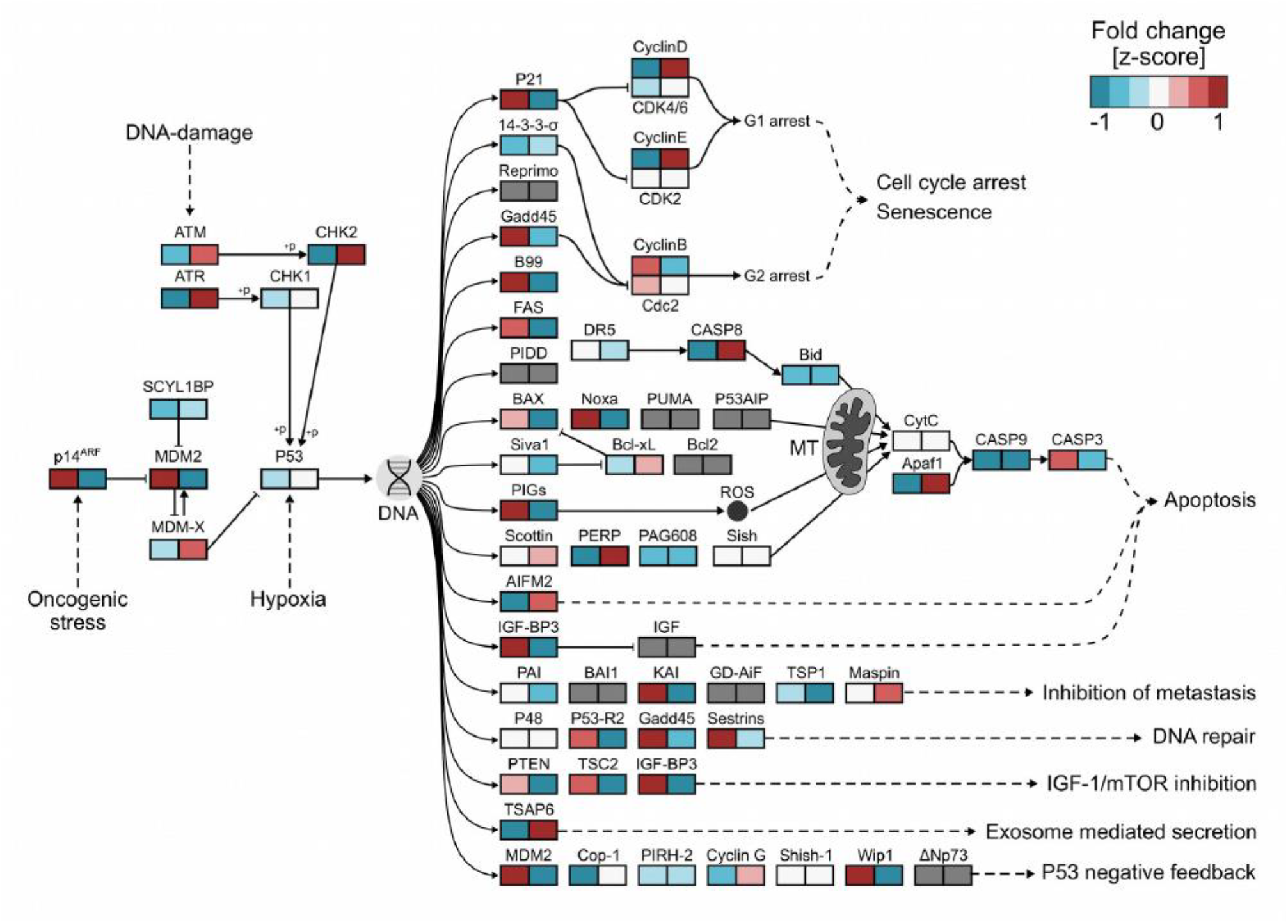
On-chip stimulation effect on P53 pathways response. Response of P53-pathway genes to dox-treatment (left panel of each gene indicates bulk fold change of doxorubicin 4h over vehicle, right panel indicates on-chip stimulation effect).

**Figure 4.**
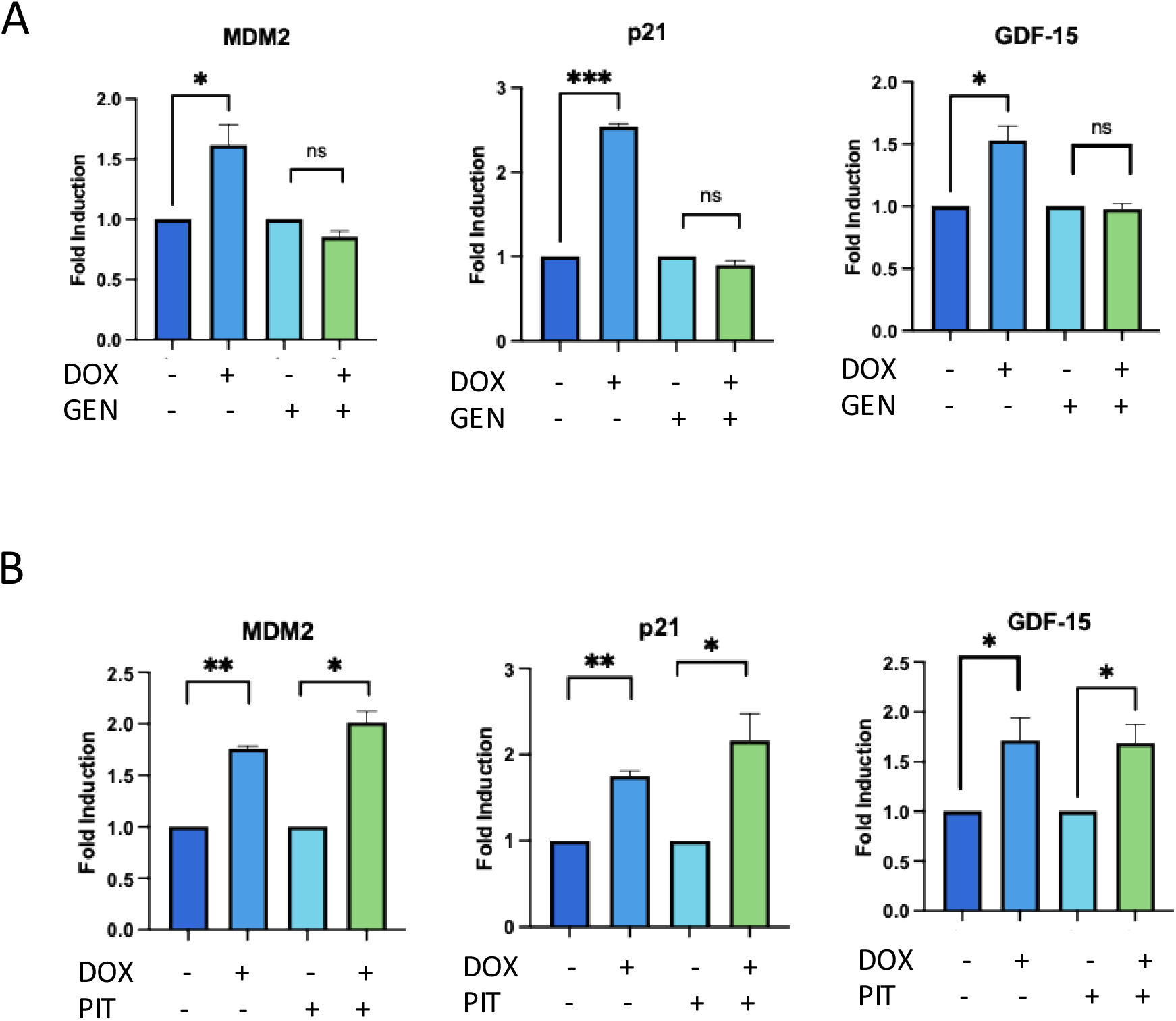
Effect on endocytosis signalling on p53 regulated genes. **(A)** Cells were treated with vehicle or doxorubicin (DOX) for 4 hours with or without Genistein (GEN), a caveolae-mediated endocytosis inhibitor, for 10 minutes. **(B)** Cell were treated with vehicle or doxorubicin (DOX) for 4 hours with or without PitStop2 (PIT), a clathrin-mediated endocytosis inhibitor, for 10 minutes. **(A)** and **(B)** RT-qPCR for MDM2, p21 and GDF-15 was performed. Values were normalized to GAPDH levels and are shown as mean ± sem. Results are from two independent experiments each performed in triplicate. *P-value < 0.05, **P-value < 0.01, ***P-value < 0.001. P-values were calculated using two-tailed Student’s t test.

Collectively, these experiments show that cell-to-cell communication is crucial to activate p53 signaling following DNA damage and that when this is impaired cells in isolation are no longer able to activate this response. The loss of activation of p53 signaling can be mimicked when cells are communicating but caveolar-mediated endocytosis is inhibited, suggesting that this is key in activating p53 response.

## Discussion

In this study, we combined microfluidic technology with scRNA-seq to investigate the DNA damage response induced by genotoxic stress in single colon cancer cells during a timecourse experiment. Furthermore, since the effect of genotoxic stress on individual cells is poorly understood, we seeded cells in sealed microfluidic chambers to explore whether preventing cell-to-cell communication following dox treatment alters gene expression regulated by DNA damage. We observed that induction of DNA-damage response marker genes including CDKN1A and MDM2 showed high variability in unstimulated cells but were coherently up-regulated following dox treatment, with reduced variability correlating with later timepoints (**Fig. S1**). Pseudotemporal ordering identified a primary trajectory corresponding to early repressed, late repressed, early activated and late activated genes by genotoxic stress (**Fig. 1B**). Interestingly, early and late repressed genes (M1 and M2) were enriched for factors modulating mitochondrial organisation and mitochondrial activity, respectively, indicating a gradual and progressive repression of mitochondrial function^1,14^. Additionally, late repressed genes were enriched for those necessary for general gene expression, including splicing and cytoplasmic translation (**Fig. 1D**). As expected, activated genes (M3 and M4) were enriched for factors directly involved in p53 signalling, including those regulating cell cycle arrest (M3). Notably, late-expressed genes (M4) were enriched for genes associated with the DDR and apoptosis^1^ (**Fig. 1D**).

Chemotherapy treatment of cancer patients with genotoxic agents such as dox preferentially kills highly proliferative tumour cells in the body^2^. However, it is unclear whether paracrine signals affect the DDR. To investigate this, we treated single HCT116 cells in bulk or isolation with dox for 4 hours, allowing or preventing cell-to-cell communication during treatment. Following this, we performed scRNA-seq to evaluate whether the absence of cell communication differentially regulates the transcriptome, accounting for covariates such as the presence of serum and low-adherent conditions in our model (**Fig. 2**). After correcting for batch effects across samples (**Fig. S1B and Methods**), we observed a clear separation between cells treated with dox in isolation (OCS) and those treated in bulk (BLK). This indicates that the absence of cell-to-cell communication has a significant transcriptional impact on the output of the DDR. However, cells in isolation remained responsive to the transcriptional changes induced by dox treatment (**Fig. 2A**). Unexpectedly, cells treated in isolation lost the ability to regulate genes associated with apoptosis, repression of MYC targets, and the p53 response (**Fig. 2B and Fig. 3**). By comparing cells treated with dox in isolation to those treated in bulk, we found that p53-modulated genes were oppositely regulated under the two conditions. For instance, while MDM2 and CDKN1A (p21) were induced by dox as expected in bulk-treated cells, they were repressed in isolated cells. Conversely, Cyclin D and CDKs, which were repressed by dox in bulk-treated cells, were more highly expressed in isolated cells compared to their bulk-treated counterparts (**Fig. 3**). Accordingly, Ingenuity Pathway Analysis (IPA) comparative analysis reveals that DNA damage, mitochondrial dysfunction, and p53 signalling are no longer enriched in cells treated with dox when cultured in isolation. This suggests that cell-to-cell communication is essential for the activation of p53. Notably, endocytosis pathways, including caveolae-mediated and clathrin-mediated endocytosis, were significantly regulated by dox (**Fig. S3**) in cells treated in bulk, but were less affected in cells treated in isolation. This suggests that, when cells are treated with dox, they must enhance endocytosis, likely to facilitate the uptake of paracrine factors necessary for the efficient modulation of p53 activity and the DDR. Using specific inhibitors of caveolae-mediated and clathrin-mediated endocytosis, we found that inhibition of caveolae-mediated endocytosis impaired the p53 response, as evidenced by the lack of activation of key p53-responsive genes (**Fig. 4**). It remains undetermined which factors are taken up by cells through caveolae-mediated endocytosis to facilitate p53 activation by dox. It has been reported that p53 itself is secreted by KRAS-mutated cells, and the p53 core domain is endocytosed by neighbouring cells via the early endocytosis pathway, directly associating with caveolin-1^15,16^. HCT116 cells harbour the KRAS^G13D^ mutation, so it is possible that p53 is internalised in these cells. However, it remains unclear why endogenous p53 appears inactive in isolated cells, suggesting that the mechanism of cell-to-cell communication modulating p53 is more complex than simple extracellular p53 internalisation. Caveolae-mediated endocytosis also facilitates the internalisation of other molecules and cargos, such as signalling molecules and exosomes. Exosomes are taken up by cells through various endocytic pathways, including caveolae-mediated endocytosis ^17^. It will be crucial to evaluate whether key activators of p53, such as MDM2, SIRT2, or miR-34 ^18^ are essential exosomal cargos responsible for this novel p53 activation process modulated by cell-to-cell communication.

## Methods

### Cell culture

HCT-116 cells were maintained in Dulbecco’s modified Eagle medium supplemented with 10% fetal calf serum, 2 mM L-glutamine, 100 U/ml penicillin, and 100 mg/ml streptomycin. Cells were treated with dox at a concentration of 0.2 μg/ml or equivalent volume of vehicle (ddh_2_0) for the indicated time. PitStop 2 (Abcam, Ab120687) was used at 10μM and Genistein (Sigma, G6649) was used at 200μM for 10 minutes and then cells were treated with dox or vehicle for 4 hours.

### Single-cell mRNA sequencing

Single-cell mRNA sequencing was performed using the Fluidigm C1 System (Fluidigm) following the manufacturer’s protocol.

#### Single-cell capture, reverse transcription and PCR

The C1 integrated fluidics circuits (IFCs) for mRNA seq (small-cell, 5–10 μm) was primed according to the standard Fluidigm mRNA-seq protocol using C1 harvest reagent, C1 preloading reagent, C1 blocking reagent, and Fluidigm cell wash buffer. Priming was performed immediately prior to sample preparation. HCT-116 cells were trypsinised and resuspended in serum-free media at a concentration of 333,000 cells/mL. The standard suspension ratio of 3:2 was used (60 µL of cell suspension was added to 40 µL of Fluidigm suspension reagent) to achieve the final cell mix for IFC loading. After mixing several time the cell preparation, 6 µL of the cell mix was loaded. Following cell loading, light microscopy using EVOS microscope, was used in order to assess each of the 96 capture sites individually and the outcome of loading was recorded in terms of the number of cells present in each capture site and whether debris were present. This information was used as a quality control step to ensure that only confirmed single cell samples were taken forward. Finally, the IFC was placed into the C1 the script “*mRNA Seq: Cell Load (1771x/1772x/1773x)*” was run. Following the loading, buffer mixes for lysis, reverse transcription and PCR were pipetted into the IFC. The IFC was then placed into the C1 system and the script *“mRNA Seq: RT & Amp (1771x/1772x/1773x)*” was run overnight. Samples were harvested from the IFC by transferring 3.5 µL of product to a 96 well plate and diluted with 10 µL C1 DNA dilution reagent. The samples corresponding to the confirmed single cell captures were highlighted according to the arrangement on the harvest plate described by Fluidigm. Samples were stored at 4ºC short term and - 20ºC long term prior to subsequent quantification and library preparation.

#### Library Preparation and sequencing

Library preparation was performed using the modified Illumina Nextera XT DNA library preparation protocol provided by Fluidigm. The cDNA concentration was quantified using the PicoGreen Assay (Quant-IT PicoGreen dsDNA Assay Kit, Thermo Fisher Scientific, P11496) using 2 µL per 96 samples processed into 2 replicates in a 384-well plate giving a final well volume of 60 µL for reading in a fluorometer. The data was then processed using the Microsoft Excel worksheet, Single-Cell mRNA Seq PicoGreen Template (Fluidigm, PN 100-6260), to quantify the library. Samples were transferred to a new 96-well plate in order to obtain a sample within the range 0.1-0.3 ng/µL by diluting with C1 Harvest Reagent. Only samples with at least 0.1 ng/µL were taken forward for library preparation. Sequencing libraries were prepared using Nextera XT DNA Library (Illumina) with a modified protocol provided by Fluidigm.

### On-chip stimulation

In order to assess the effect of dox on cell-to-cell communication, a modified C_1_ workflow developed by Shalek and colleagues^13^ was used. Specifically, unstimulated HCT116 cells were loaded on the C1 chip and microscopy of each capture site performed to ensure single cell capture and viability. After that the cells were stimulated on chip with media containing dox or vehicle for 4 hours. Following incubation, samples were processed for scRNA-seq as previously described.

### RNA isolation and RT-qPCR assays

Total RNA from cultured cells was extracted using TRI Reagent (Sigma) followed by Direct-zol RNA Miniprep (Zymo Research) following the manufacturer’s instructions including DNase I treatment. cDNA was synthesized from 1 μg of purified DNase-treated RNA using RevertAidTM M-MuLV reverse transcriptase and random hexamer primers (Thermo Scientific), according to the manufacturer’s protocols. We performed qRT-PCR on a StepOne™ Real-Time PCR System using Fast SYBR® Green Master Mix (both from Applied Biosystems). The primer sequences are as follow: GAPDH forward primer 5’-TGAAGGTCGGAGTCAACGGATTT-3’ and reverse 5’-GCCATGGAATTTGCCATGGGTGG-3’; MDM2 forward primer 5’-CCATTGAACCTTGTGTGATTTGTC-3’ and reverse 5’-TTCCTTTTCTTTAGCTTCTTTGCAC-3’; GDF-15 forward primer5’-TCTCAGATGCTCCTGGTGTTG-3’ and reverse 5’-TTCCGCAACTCTCGGAATCT-3’.

### Pathway Analyses

Gene set enrichment analysis was performed in R using gene log2 fold change values as input metrics using the clusterProfiler package v3.12. and Comparison pathway analysis was performed using Qiagen Ingenuity Pathway Analysis (IPA) (https://digitalinsights.qiagen.com/products-overview/discovery-insights-portfolio/analysis-and-visualization/qiagen-ipa/. Two datasets were used for the comparison: 1) the DOX vs vehicle in bulk condition and 2) the DOX vs vehicle in on-chip stimulation. Data was visualised using the – log (p-value).

### Single cell RNA-seq analysis

Reads were aligned to the transcriptome (GRCh38 Gencode v26) supplemented with ERCC spike in sequences utilizing STAR v2.5.0 with the following non-default parameters: --outFilterType BySJout --outFilterMultimapNmax 20 -- alignSJoverhangMin 8 --alignSJDBoverhangMin 1 –outFilterMismatchNoverLmax 0.04 --alignIntronMin 20 --alignIntronMax 1000000 --alignMatesGapMax 1000000. Cells were pre-filtered for total detected features (>4k) and fraction of reads aligned to control features (ERCC + mitochondrial) (<0.7). Batch correction and normalization performed with RUVseq RUVs^19^, with number of unwanted variance factors k=10 as estimated by PCA analysis. Cell cycle phase was predicted with the cyclone pretrained classifier as implemented in scran v1.4.0^20^ using standard human reference features^21^. Pseudotemporal trajectory inference was performed using SCORPIUS^22^ on normalized expression data reduced to 3 dimensions by eigen analysis of the dissimilarity matrix using spearman distances, k=4 clusters and default parameters for curve fitting.

## Data availability

Raw and processed data has been deposited to GEO accession number GSE298355.

## Author Contributions

L.C., S.O. and J.S. conceived and supervised the study. M.R. conducted all bioinformatic analyses. A.D. and N.F. conducted experiments. L.C. and S.O. designed and conducted single-cell RNA-seq experiments, and wrote the paper. J.S. supplied reagents and edited the paper. All authors approved the final draft of the work.

## Acknowledgements

This study was funded by Action Against Cancer (AAC). In particular, the Colin McDavid Family Trust, Alessandro Dusi, Mike Hunter, Kevin and Noreen Coyle and Cheryl Whitehead, BHM. The authors would also like to thank the London Institute of Medical Sciences Genomics Facility and Imperial BRC Genomics Facility for technical assistance during the single-cell protocol and sequencing.

## Declaration of interests

J.S. declares his conflicts at: https://www.nature.com/onc/editors, and none are relevant here. The remaining authors declare no competing interests.

