## Supplementary Figures for "Single-cell transcriptomics uncovers that cell-to-cell communication controls p53 activation"

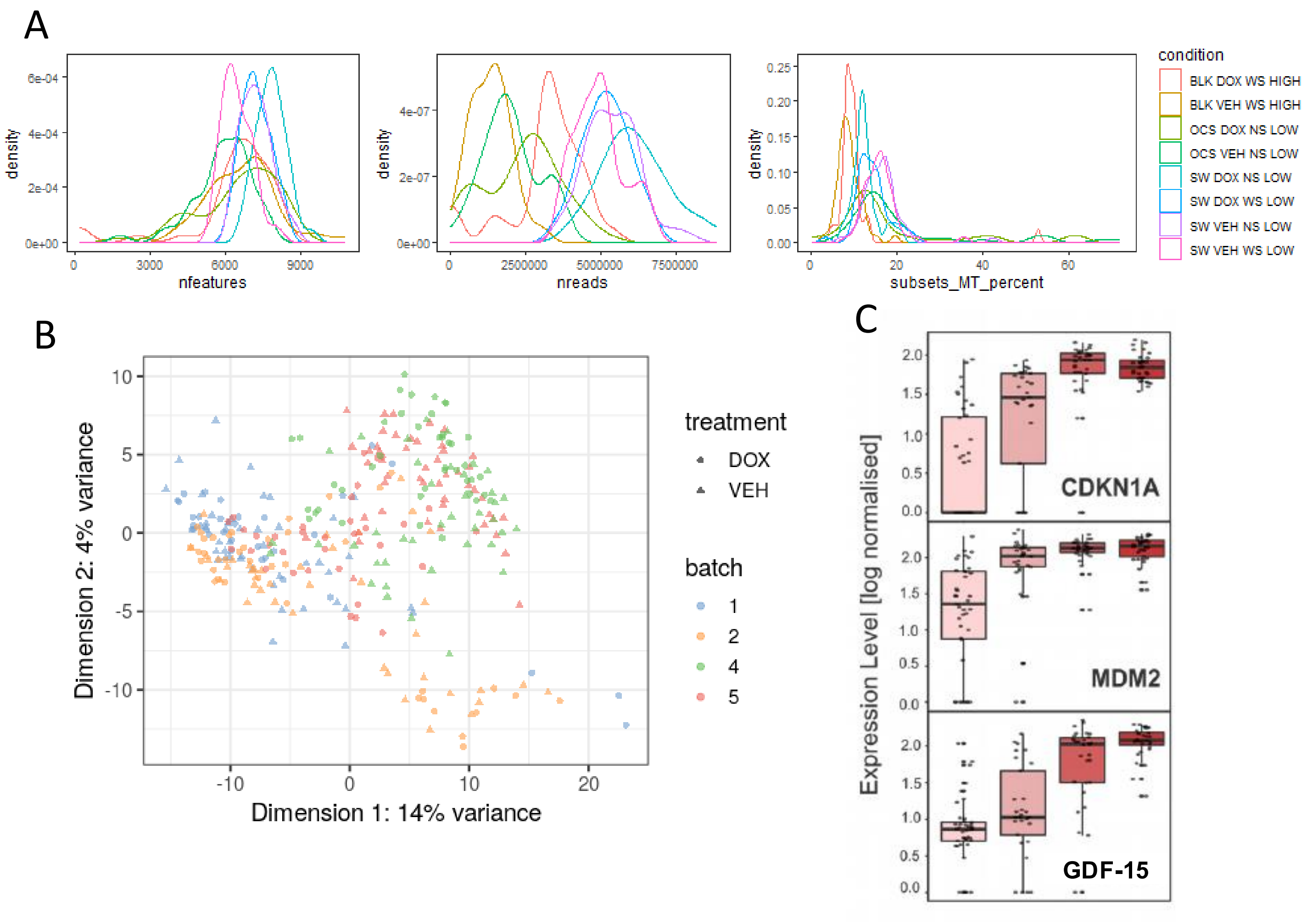


**Supplementary Figure 1. A)** Distribution of QC metrics for cells post-filtering. **B)** PCA of bulk-treated samples prior to batch correction demonstrates clustering by batch, not treatment condition. **C)** Key DNA-damage response genes including CDKN1A and MDM2 show expected pattern of up-regulation in response to dox treatment.

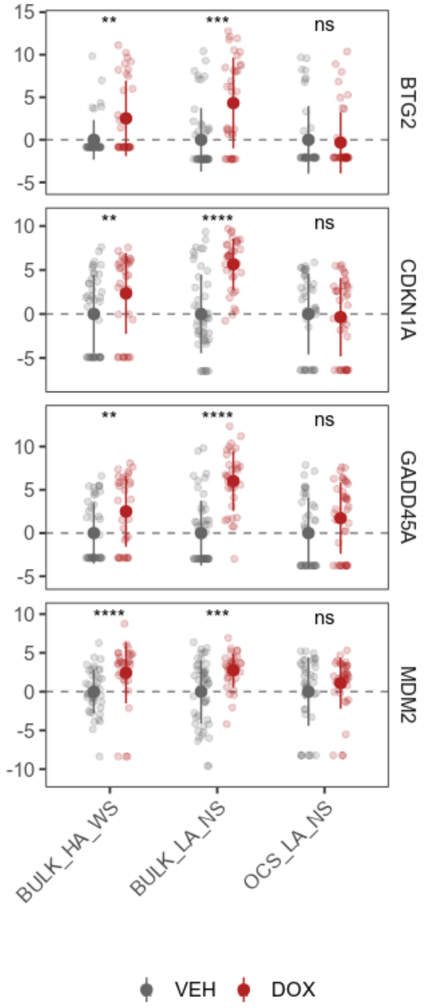


**Supplementary Figure 2.** Expression of key p53 response genes in bulk conditions (adherence with serum BULK_HA_WS, low adherence and no serum BUL_LA_NS) vs on-chip stimulation (low adherence and no serum OCS_LA_NS)

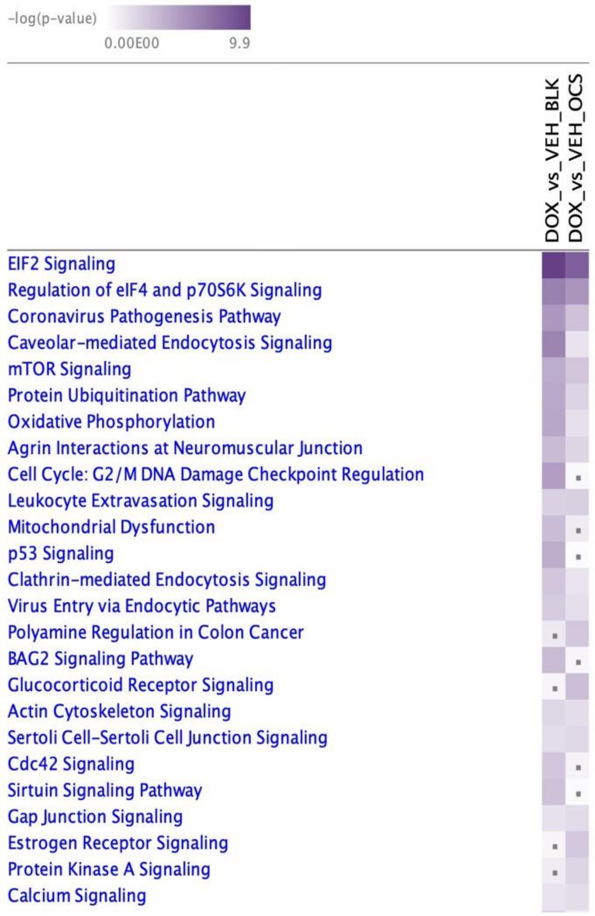


**Supplementary Figure 3.** Ingenuity Pathway Analysis (IPA) comparison analysis shows that DNA damage, mitochondrial dysfunction and p53 signalling are no longer enriched in on chip stimulation (OCS). Dots indicate the pathways that are below the – log (p-value) cutoff value of 1.3.
